# Effects of water decontamination methods and bedding material on the gut microbiota

**DOI:** 10.1101/325878

**Authors:** Willie A. Bidot, Aaron C. Ericsson, Craig L. Franklin

**Affiliations:** Comparative Medicine Program, University of Missouri, Columbia, Missouri, United States of America; University of Missouri Metagenomics Center, University of Missouri, Columbia, Missouri, United States of America; Mutant Mouse Resource & Research Center, University of Missouri, Columbia, Missouri, United States of America

**Author notes:** Corresponding author, (CF).

## Abstract

Rodent models are invaluable to understanding health and disease in many areas of biomedical research. Unfortunately, many models suffer from lack of phenotype reproducibility. Our laboratory has shown that differences in gut microbiota (GM) can modulate phenotypes of models of colon cancer and inflammatory bowel disease. We and others have also shown that a number of factors associated with rodent research, including vendor, cage system, and bedding can alter GM. The objective of this study was to expand these studies to examine the effect of additional bedding materials and methods of water decontamination on GM diversity and composition. To this end, Crl:CD1 (ICR) mice were housed on corn cob or compressed paper chip bedding and provided water that was decontaminated by four commonly used procedures: reverse osmosis, autoclaving, sulfuric acid treatment, or hydrochloric acid treatment. Feces was collected at day 0, and at day 28 (endpoint), fecal and cecal samples were collected. DNA was extracted from samples, amplified by PCR using conserved bacterial primer sets and subjected to next generation sequencing. Sequence data were analyzed using Qiime and groups were compared using principal coordinate analysis (PCoA) and permutational multivariate analysis of variance (PERMANOVA). Two factor PERMANOVA of cecal GM data revealed significant changes when comparing bedding and water decontamination methods, while no significant effects were noted in the fecal GM data. Subsequent PERMANOVA and PCoA of cecal data revealed that several combinations of bedding and water decontamination methods resulted in differing GM, highlighting the complexity by which environmental factors interact to modulate GM.

## Introduction

In recent years there has been a substantial increase in studies focusing on the microorganisms present in the gastrointestinal tract (GIT). The gut microbiota (GM) is known to play crucial roles in digestion, immune status development, and pathogen resistance, and differences in GM have been associated with differences in health and disease susceptibility (1–3). Certain characteristics in the GM have been associated with diseases of the GIT (4), as well as diseases in other body systems such as the central nervous system (5, 6). Rodent models have emerged as a highly valuable tool to determine the role of the GM in both health and disease. Several studies have demonstrated that the highly dynamic GM is influenced by a variety of environmental factors, and can in turn impact rodent model phenotypes (7). Recently, the use of mouse models has been questioned due to the lack of reproducibility (8). These limitations have spurred efforts from several institutions such as the National Institutes of Health (NIH) to improve reproducibility of animal research (9). Our laboratory has focused on the microbial composition of the GIT as an important contributing factor in phenotypic variability of rodent disease models (10, 11). We previously found that the GM differs depending on the source and genetic background of the mouse (12). Even mice of genetically similar backgrounds from the same producer can have differing GM depending on the institution in which they are housed, suggesting that the environment is a major factor in the determination of the GM (13). We have also demonstrated significant changes in the GM in response to housing conditions (14), further corroborating the importance of environmental factors in shaping the GM. Given that the GM significantly impacts model phenotypes, these data substantiate the need to consider how different husbandry factors may influence the GM.

Husbandry factors such as light cycle, temperature, bedding, and handling can be seen as subtle factors that can affect the outcome of rodent experiments (7, 15). Factors such as temperature and light cycle can be controlled by proper building maintenance. Bedding, a factor that is often overlooked, can greatly differ between facilities.

Another factor that can be overlooked is the water that is offered to rodents. Several water decontamination methods are commonplace in contemporary rodent facilities. Methods include filtering the water to physically remove contaminants (e.g., reverse osmosis), or procedures to kill bacteria (e.g., UV light or acidification). Differing water decontamination methods have been shown to impact model phenotypes. For example, the low pH of acidified water was associated with phenotype changes in a mouse diabetic model (16, 17). Moreover, water chlorination, when compared to tap water alone was shown to change the phenotype of a mouse model of colorectal cancer (18). It is unclear whether different water decontamination methods influence the GM, and in turn, become a potential experimental variable that contributes to inadequate reproducibility of model phenotypes.

Bedding is another component of husbandry that varies between animal facilities. Corn cob bedding is one of the most commonly used bedding materials because of its high absorbency (19) and low cost; however other paper-based bedding materials are becoming popular. Differing bedding materials have also been linked to model phenotype changes and changes in the GM (20, 21), but controlled studies have yet to be performed. To address how water decontamination and bedding shape the GM, we exposed mice to water decontaminated by four different methods: autoclaving with reverse osmosis (RO), autoclaving with hydrochloric acid (HCl), autoclaving with sulfuric acid (H_2_SO_4_), and autoclaving alone (Autoclaved). We also exposed the mice to two different bedding materials, corn cob and paperchip, and evaluated the interaction between water and bedding as drivers of GM composition change. Composition of the GM was determined by targeted amplicon sequencing using DNA extracted from feces and cecal content. Samples were collected upon arrival and four weeks after being exposed to either of the water and bedding combinations. Robust statistical methods then were used to determine main effects of, and interaction between water and bedding.

Understanding how husbandry factors, such as water decontamination or choice of bedding, can influence the GM is a critical first step toward improving reproducibility in animal models.

## Materials and Methods

### Ethics statement

This study was performed in accordance with the recommendations put forth in the Guide for the Care and Use of Laboratory Animals and were approved by the University of Missouri Institutional Animal Care and Use Committee (MU ACUC protocol #8720).

### Survey

A water decontamination survey (S1 Table) was sent out on December 15, 2015 using the Compmed listserv (CompMed, AALAS, Memphis, TN), a listserv that is used for discussion of subjects of comparative medicine, laboratory animals, and topics related to biomedical research.

On March 21, 2016 a question survey about bedding was sent using the same email list. The survey included the following; “I am conducting a survey on the different types of bedding used in rodent facilities. Please send me a response with the type of bedding used in your facility. Specifics on the bedding will be appreciated (size, form, etc.).” Results from both surveys were recorded until the last response was received on March 29, 2016.

### Mice

Six to eight week-old female outbred Crl:CD1 (ICR) mice (*n* = 96) were purchased from Charles River Laboratories (Wilmington, MA) in a single order, and housed in the same room and maintained under barrier conditions in microisolator cages on individually ventilated cage-racks (Thoren, Hazleton, PA), filled with either compressed paper (Paperchip^®^ Brand Laboratory Animal Bedding, Shepherd Specialty Papers, Watertown, TN) or corn cob bedding (Bed-o’Cobs 1/8″, The Andersons Inc., Maumee, OH), with *ad libitum* access to autoclaved rodent chow (LabDiet 5008 Purina, St. Louis, MO) and water, under a 14:10 light/dark cycle. The cages contained a nestlet for enrichment and 4 mice per cage. The water offered was municipal water which was decontaminated using the methods explained below. Using a random number generator, mice were randomly assigned to one of the water and bedding combinations. Mice were determined to be free of overt and opportunist bacterial pathogens including *Citrobacter rodentium, Clostridium piliforme, Corynebacterium kutscheri, Helicobacter* spp., *Mycoplasma* spp., *Pasteurella pneumotropica, Pseudomonas aeruginosa, Salmonella* spp., *Streptococcus pneumoniae*; MHV, MVM, MPV, MNV, TMEV, EDIM, LCMV, MAV1, MAV2, Polyomavirus, PVM, REO3, Ectromelia virus, and Sendai virus; intestinal protozoa including *Spironucleus muris, Giardia muris, Entamoeba muris,* trichomonads, and other large intestinal flagellates and amoeba; intestinal parasites including pinworms and tapeworms; and external parasites including all species of lice and mites, via quarterly sentinel testing performed by IDEXX BioResearch (Columbia, MO). At the end of study, mice were humanely euthanized via inhaled carbon dioxide, in accordance with the AVMA Guidelines for the Euthanasia of Animals: 2013 Edition, followed by cervical dislocation as a secondary means.

### Water decontamination methods

The four water decontamination methods selected were tap water (autoclaved), reverse osmosis, and acidification with either hydrochloric acid or sulfuric acid. Filled water bottles from all four groups were autoclaved prior to use. Reverse osmosis filtration used a Milli-Q^®^ Direct system (Merck KGaA, Darmstadt, Germany). Acidification was performed using an automated bottle filler which titrated the water with sulfuric acid (model 9WEF, Tecniplast, Buguggiate, Italy) or hydrochloric acid (model Basil 1100, Steris Corporation, Mentor, OH) to a target pH of 2.5 (range 2.3 to 2.7). Water pH was verified using a handheld pH meter (pHTestr^®^ 10, Oakton Instruments, Vernon Hills, IL).

### Sample collection

Freshly evacuated fecal pellets were obtained from each mouse on the day of arrival. These samples were collected by transferring each mouse to a separate clean microisolator cage containing no bedding, and allowing the mouse to defecate normally. Fecal pellets were then collected with a sterile wooden toothpick. At day 28 post-arrival mice were humanely euthanized and cecal and fecal samples were collected using aseptic technique. Briefly, each region of the gut was exteriorized to allow collection of samples from roughly the same site of each animal. Cecal samples comprised the entire cecal contents; and fecal samples represented the most distal fecal bolus present in the colon, excluding boli within the rectum proper. Instruments used for collection were flamed and allowed to cool between all samples. Following collection, all samples were placed in 2 mL round-bottom tubes containing a 0.5 cm diameter stainless steel bead. All samples were stored in a −80°C freezer until extraction was performed.

For DNA extraction, 800 μL of lysis buffer (22) was added to each tube containing the sample and then mechanically disrupted using a TissueLyser II (Qiagen, Venlo, Netherlands). After mechanical disruption, tubes were incubated at 70°C for 20 minutes with periodic vortexing. Samples were then centrifuged at 5000×g for five minutes at room temperature, and the supernatant transferred to a clean 1.5 mL Eppendorf tube. Two hundred μL of 10 mM ammonium acetate was added to lysates, mixed thoroughly, incubated on ice for five minutes, and then centrifuged as above. Supernatant was then mixed thoroughly with one volume of chilled isopropanol and incubated on ice for 30 minutes. Samples were then centrifuged at 16000×g for 15 minutes at 4°C. The supernatant was aspirated and discarded, and the DNA pellet washed several times with 70% ethanol and resuspended in 150 μL of Tris-EDTA. 15 μL of proteinase-K and 200 μL of Buffer AL (DNeasy Blood and Tissue kit, Qiagen) were added and samples were incubated at 70°C for 10 minutes. 200 μL of 100% ethanol was added and the contents of each tube were transferred to a spin column from the DNeasy kit. DNA was then purified according to the manufacturer’s instructions and eluted in 200 μL of EB buffer (Qiagen). Purity of DNA was assessed via spectrophotometry (Nanodrop, Thermo Fisher Scientific, Waltham, MA); yield was determined via fluorometry (Qubit, Life Technologies, Carlsbad, CA) using quant-iT BR dsDNA reagent kit (Invitrogen, Carlsbad, CA).

### 16S rRNA library preparation and sequencing

The extracted DNA was amplified and sequenced at the University of Missouri DNA Core facility, as previously described (23). Briefly, an amplicon library of the V4 region of the 16S rRNA gene was generated using normalized DNA as a template. Using single-indexed universal primers (U515F/806R) flanked by Illumina standard adapter sequences with PCR parameters of 98°C^(3 m)^ + [98°C^(15 s)^ + 50°C^(30 s)^ + 72°C^(30 s)^] × 25 cycles + 72°C^(7 m)^. Amplicons were then pooled for sequencing using Illumina MiSeq and V2 chemistry with 2×250 bp paired-end reads.

### Informatics

All assembly, filtering, binning, and annotation of contiguous sequences was performed at the University of Missouri Informatics Research Core Facility, as previously described (23), with one exception: Operational taxonomic units (OTUs) were annotated using BLAST (24) against the SILVA database (25) of 16S rRNA sequences and taxonomy rather than the Greengenes database used in our previous studies.

### Statistical Analysis

Samples receiving less than 10,000 sequence reads were omitted from analysis. Differences in beta-diversity of all groups were tested via a two-way and one-way PERMANOVA of ranked Bray-Curtis (shared abundances of OTUs) similarity index using the open access Past 3.14 software package (26). Principal coordinate analysis (PCoA) was also performed using the Past software package and the relative abundance data was fourth-root transformed to normalize the data. OTU richness and diversity indices were tested for normality using the Shapiro-Wilk method; differences were then tested via two-way ANOVA for normal data or Kruskal-Wallis ANOVA on ranks for non-normal data using SigmaPlot 12.3 (Systat Software Inc., San Jose, CA). Significant OTUs, hierarchical clustering, and random forest analysis were performed using cube root-transformed sequence data using open access MetaboAnalyst 3.0 (http://www.metaboanalyst.ca) (27). We considered *p* values less than 0.05 significant.

## Data availability

All reported data have been deposited in the National Center for Biotechnology Information (NCBI) Sequence Read Archive (SRA) under BioProject accession number PRJNA453789.

## Results

### Water decontamination methods and bedding material survey results

To determine what methods are being used in contemporary housing, two surveys were conducted. A survey was conducted to identify water decontamination methods that are available and being used across different rodent facilities (Fig 1A). A total of 39 responses from 19 institutions were received and 13 different water decontamination methods were identified. The most common method used was reverse osmosis (RO) followed by acidification with hydrochloric acid. Autoclaving tap water was the third most common method and is a more popular practice in smaller rodent facilities. Acidifying the water as a whole is a very common practice, with a total of 10 facilities acidifying their water and three different acidifiers identified; hydrochloric, sulfuric, and phosphoric acid. Due to barrier restrictions within the vivarium in which these studies were conducted, water received by all groups was autoclaved, while certain mice received water that was also purified via RO or acidified via HCl or H_2_SO_4_.

**Fig 1.**
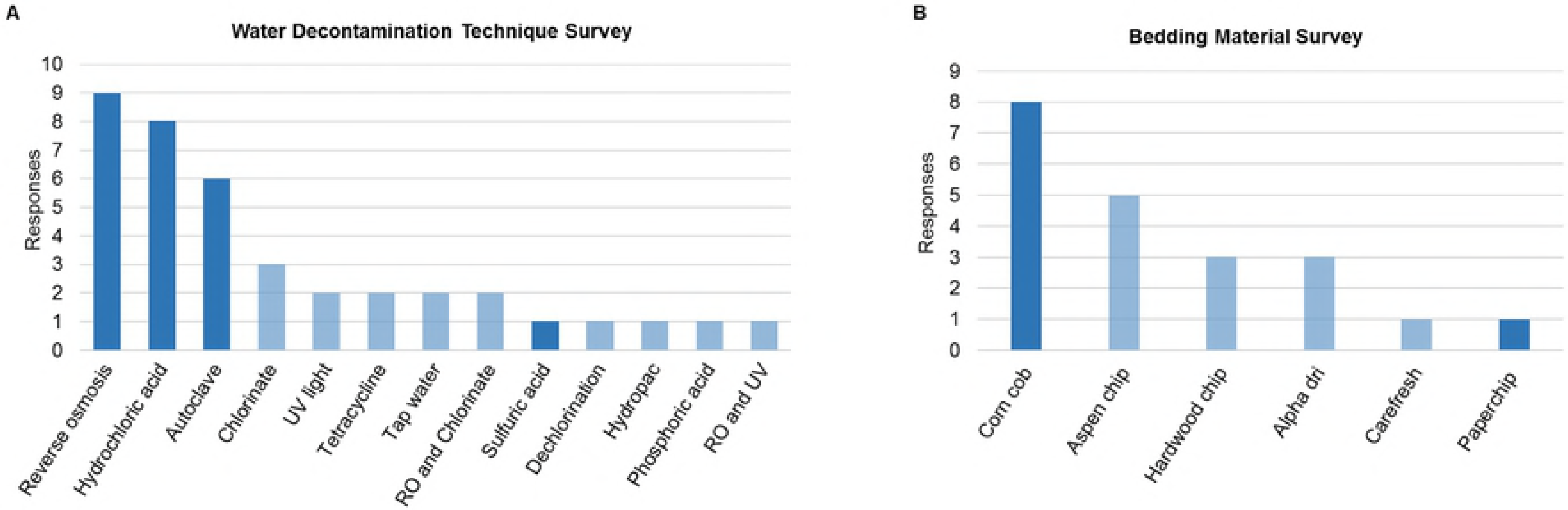
Water decontamination technique and bedding material surveys. Results of surveys on different water decontamination techniques performed on December 2015 (A), and survey on different bedding materials performed on March 2016 (B). Surveys were performed through Compmed listserv. Darkened bars represent water and bedding used in this study.

A second survey was conducted to identify what bedding materials are being used in different rodent facilities (Fig 1B). A total of 11 institutions responded, with a total of 21 responses. Corn cob bedding was the most common bedding material used (8/21), followed by aspen chip (5/21). Three different paper-based beddings were identified; alpha dri (3/21), carefresh (1/21), and paperchip (1/21). Based on availability at our institution, corn cob and paperchip were used, and eight groups of mice (2 beddings × 4 water treatments) were established in a fully crossed study design.

### Main effects of water decontamination methods and bedding material

When subjectively evaluating the composition of the fecal samples, it is difficult to observe distinct differences in relative abundance of OTUs (S1 Fig A). In contrast, subjective evaluation of the composition of cecal samples revealed a water-treatment dependent pattern in the most abundant OTUs such as UC Family Bacteroidales S24-7 1 and Lachnospiraceae NK4A136 group sp. 2 (S1 Fig B). For evaluation of the overall main effects of water decontamination methods and bedding material on the GM composition, a two-way PERMANOVA of ranked Bray-Curtis similarity index was performed (Table 1). Surprisingly, there were no significant differences in the microbiota composition of the fecal samples (FM, fecal microbiota). However, when cecal samples (CM, cecal microbiota) were examined, significant main effects were detected in both water decontamination methods (*p*=0.001) and bedding material (*p*=0.023). To evaluate all of the different water and bedding combinations, a one-way PERMANOVA with pairwise comparison of cecal communities was performed (Table 2). Out of 28 comparisons, 16 (57%) were significantly different demonstrating the complexity by which water decontamination and bedding interactions can influence microbiota.

**Table 1.** Main effects of bedding and water treatment on the fecal and cecal microbiota.

| Endpoint Data<br>Two-way Permutational Multivariate Analysis of Variance (PERMANOVA) |  |  |  |  |
| --- | --- | --- | --- | --- |
| Source | Fecal |  | Cecal |  |
| | F | $p$ | F | $p$ |
| Bedding | 1.74 | 0.117 | 2.52 | 0.023 |
| Water Decontamination Method | 0.712 | 0.626 | 4.16 | <0.001 |
| Interaction | -0.500 | 0.095 | 0.24 | 0.106 |

Two-way PERMANOVA of ranked Bray-Curtis similarity indices results from endpoint (Day 28) fecal and cecal samples. Considered $p<0.05$ significant.

**Table 2.**
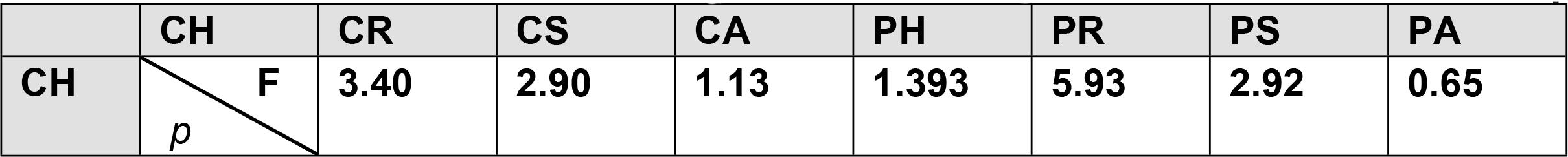
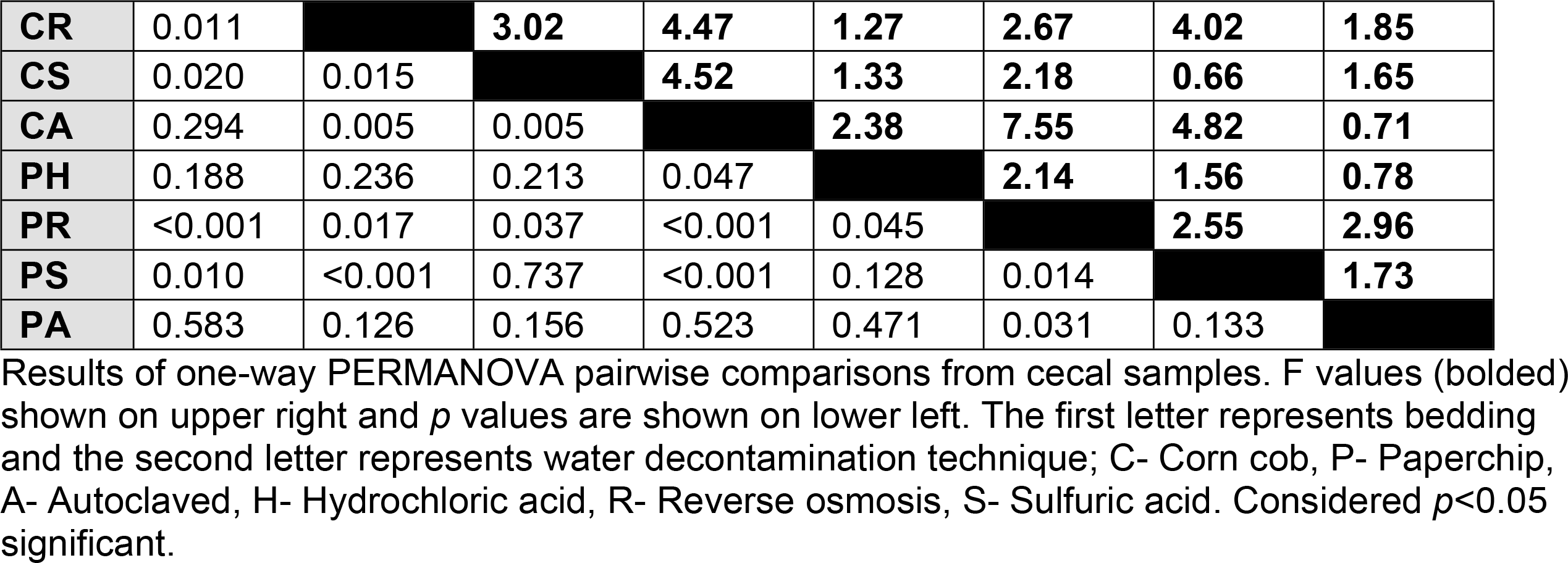
Pairwise PERMANOVA of Bray-Curtis similarity indices between cecal samples.

Differences in CM composition between groups were also visualized by PCoA. All comparisons except for one revealed separation of groups with some overlap in PCo1 vs PCo2. Fig 2 shows selected comparisons and further emphasizes the complexity of interaction between the two factors examined.

**Fig 2.**
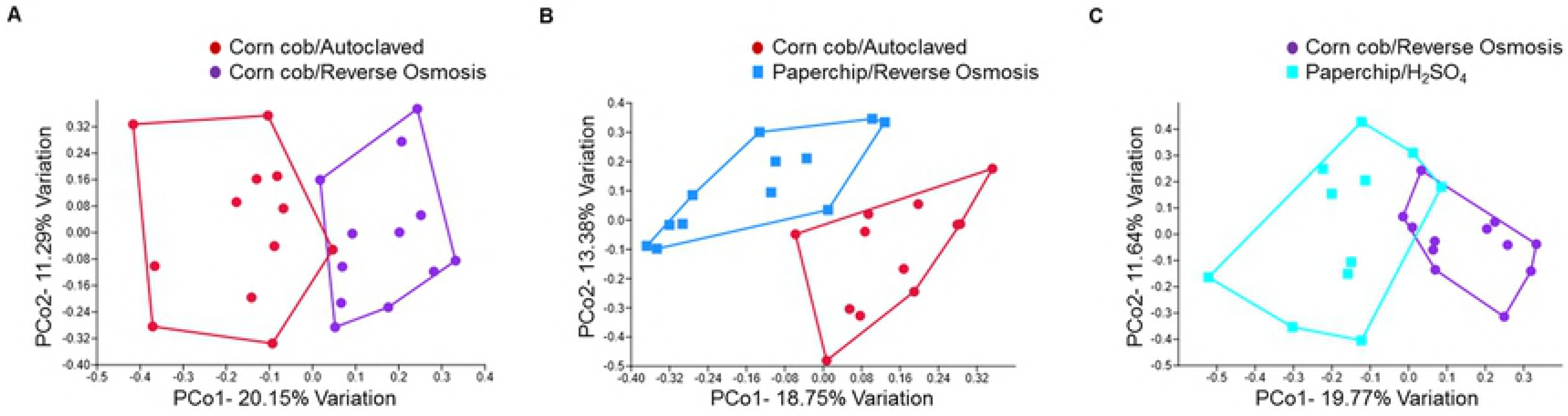
Principal Coordinate Analysis (PCoA) of pairwise comparisons of selected groups. PCoAs of comparisons between groups housed in corn cob bedding offered either reverse osmosis or autoclaved treated water (A), paperchip or corn cob bedding offered reverse osmosis or autoclaved treated water respectively (B), and corn cob or paperchip bedding offered reverse osmosis or H2SO4 treated water respectively (C). Significant differences (p<0.05) in cecal microbiota composition between these group comparisons were detected via pairwise PERMANOVA (Table 2).

### Differences in richness and alpha diversity

To measure richness of each group the number of distinct OTUs was counted for each sample. In both FM and CM there was a significant main effect of bedding on richness (S2 and S3 Tables). Overall there was a significant decrease in number of OTUs (i.e., richness) in mice housed on paperchip bedding when compared to corn cob (Fig 3 A and B). In the FM, some individual significant pairwise comparisons were demonstrated in groups offered the same water source but different beddings; HCl, RO, and H_2_SO_4_ (Fig 3A). In the CM, there was also a significant main effect of water (S3 Table), and several pairwise comparisons were significantly different (Fig 3B). Within each bedding type, samples from mice receiving autoclaved water with no additional treatment had the lowest richness, suggesting that all additional treatment methods are associated with increased richness.

**Fig 3.**
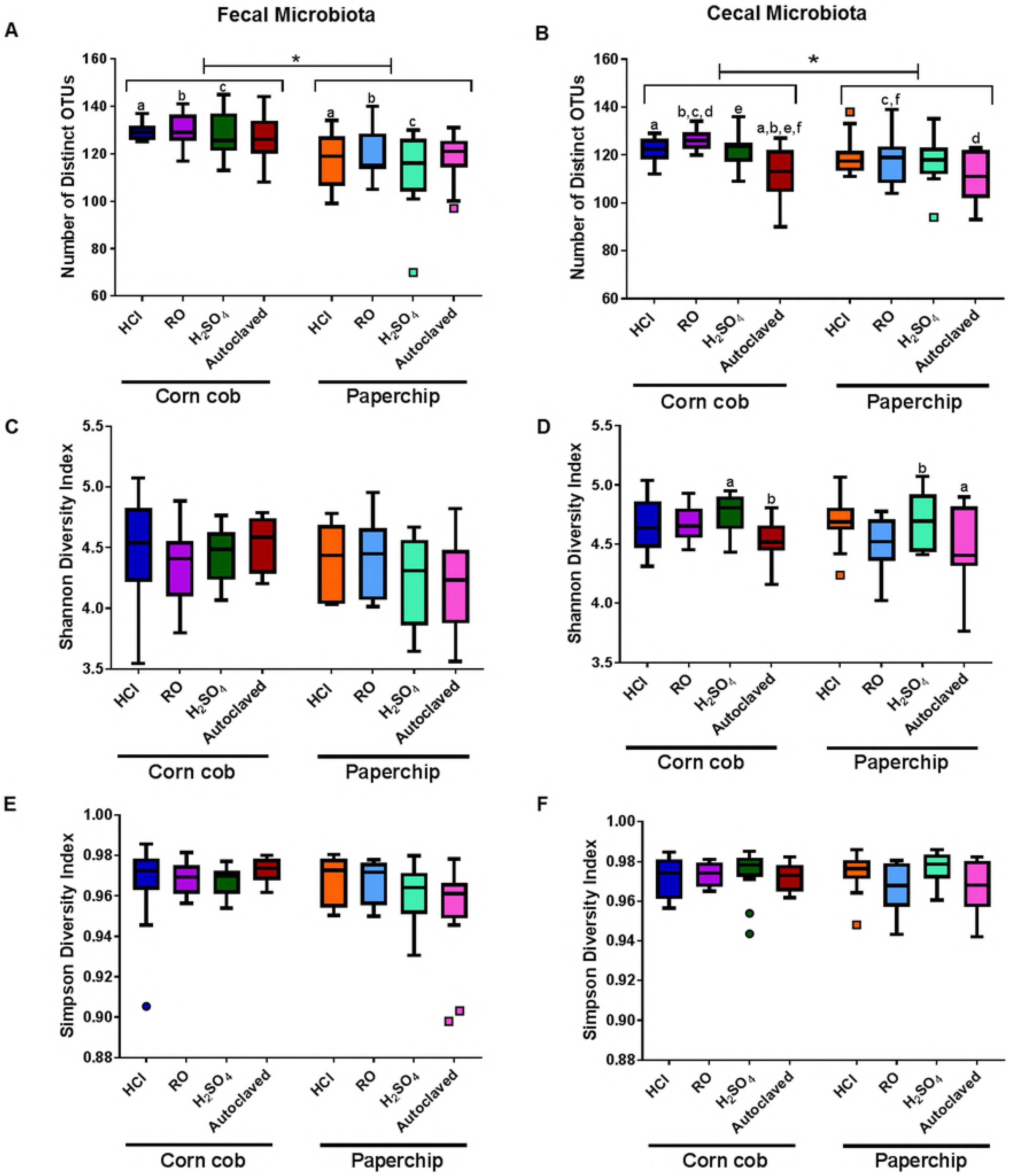
Richness and diversity of fecal and cecal microbiota at endpoint. Tukey’s box plot of endpoint fecal and cecal Richness (number of distinct OTUs) (A and B), Shannon Diversity Index (C and D), and Simpson diversity index (E and F). Like letters indicate significantly (p<0.05) different comparisons via two-way ANOVA followed by Tukey’s post hoc test. Bottom axis represents combination of bedding and water decontamination method: autoclaving with reverse osmosis (RO), autoclaving with hydrochloric acid (HCl), autoclaving with sulfuric acid (H2SO4), and autoclaving alone (Autoclaved).

Alpha diversity of the samples was calculated using the Shannon and Simpson diversity indices, which combine richness and evenness of the OTUs. No significant differences were observed in the FM in either of the diversity indices (S2 Table). In the CM however, there was a significant effect of water decontamination method (S3 Table) on α-diversity, as measured via the Shannon index. In pairwise comparisons, several significant differences were demonstrated between groups housed with autoclaved or H_2_SO_4_-treated water and housed in paper or corn cob bedding (Fig 3D). That said, no significant differences were observed in Simpson diversity index of CM (Fig 3F). Collectively, these data indicate that both bedding and water treatment methods primary influence the richness, but not the distribution of the CM, and that the bedding dependent effects on richness are maintained in the FM.

### Variation in OTU abundances and group clustering

A hierarchical cluster analyses was performed to demonstrate how individuals within experimental groups clustered according to the relative abundance of the 25 most variable OTUs as determined by ANOVA (S2 Fig). All cecal samples were represented in the analysis and were classified by treatment group. As in a PCoA, samples with similar composition cluster more closely to each other. Based on these most variable taxa, samples from several treatment groups clustered together loosely. Specifically, samples from the majority of mice receiving autoclaved water were grouped on one distal arm of the dendogram, while samples from mice receiving RO-treated water formed a separate distinct branch.

We also performed a Random Forest (RF) analysis as a means to predict OTUs that were preferentially influenced by the different husbandry conditions (Fig 4). The analysis selected 15 OTUs as important classifiers for the different husbandry conditions. Additionally, a two-way ANOVA was performed to examine which factor influenced the relative abundance most and compare to the RF results. Fourteen of the 15 OTUs selected by the RF were significantly different in the two-way ANOVA (S4 Table). These OTUs (Family XIII UCG-001 sp., *Lachnoclostridium* sp. 1, UC Family *Clostridiales* vadinBB60 group 1, *Akkermansia sp., Ruminococcaceae* UCG-009 sp., *Ruminiclostridium* 5 sp. 4, UC Family *Clostridiales* vadinBB60 group 3, UC Family *Peptococcaceae* 1, *Ruminiclostridium* 5 sp. 2, *Shuttleworthia* sp., *Lactobacillus* sp., *Enterorhabdus* sp., UC Order *Mollicutes* RF9 1, and *Anaerostipes* sp.) were detected in both analyses and thus represent candidate taxa most influenced by the water/bedding combination. All OTUs determined to be different via ANOVA demonstrated a main effect of water-treatment (S4 Table). The water main effect was visualized in several OTUs with the RF analysis (Fig 4). For example, Family XIII UCG-001, *Akkermansia* sp., UC Family *Peptococcaceae* 1, and *Anaerostipes* were all most abundant in both groups receiving autoclaved water. *Enterorhabdus* was more abundant in both H_2_SO_4_ water groups, and less abundant in the RO groups. *Shuttleworthia* was more abundant in both H_2_SO_4_ water groups, while least abundant in the autoclaved groups.

**Fig 4.**
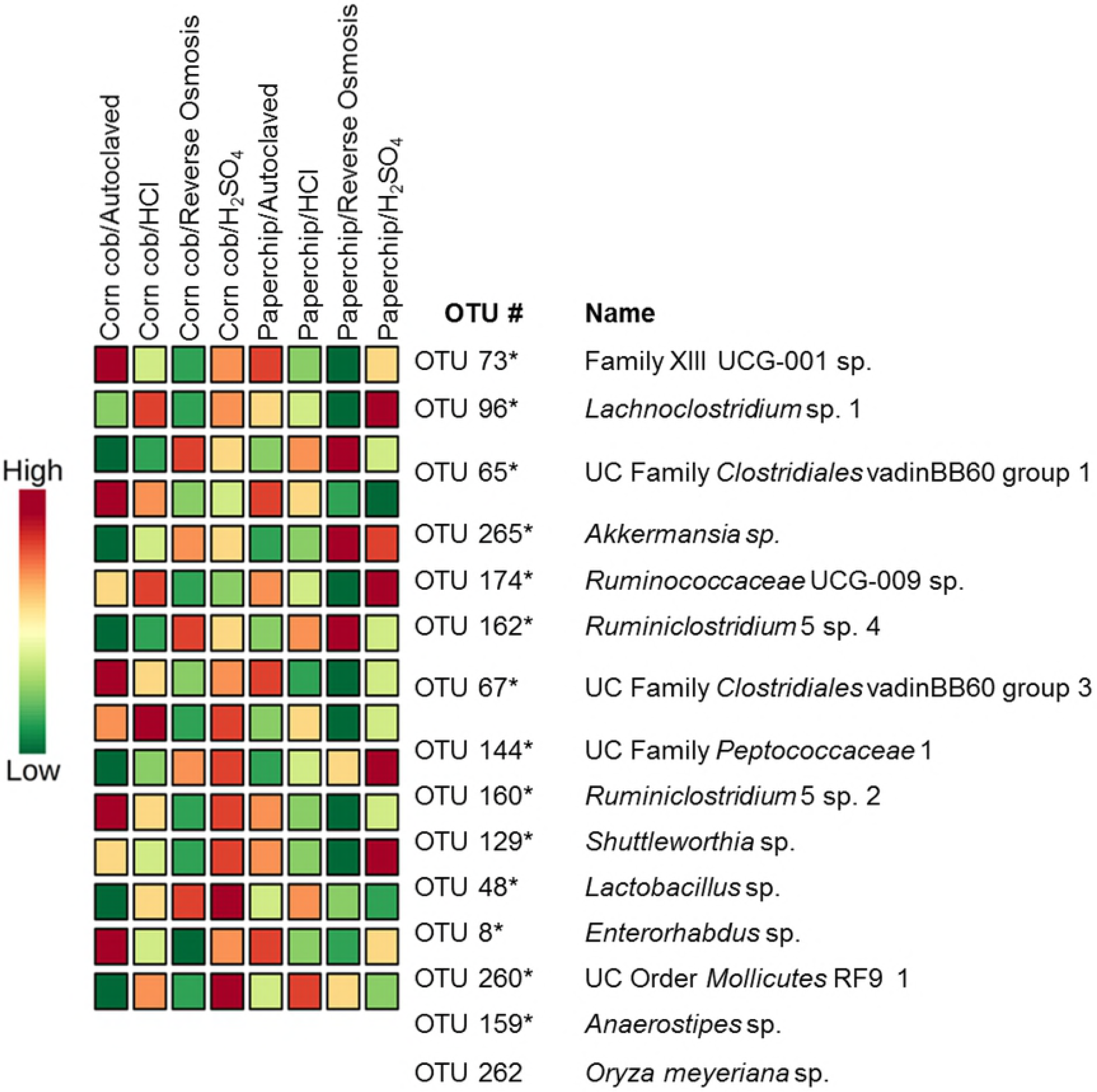
Random forest analysis selection of operational taxonomic units (OTUs). Random forest analysis of the selected most important OTUs to classify groups. Scale on left represents abundance of OTUs in each group. Asterisks represent OTUs that were significant in two-way ANOVA (S3 Table).

Collectively, the data presented above show that bedding material and water decontamination methods can influence the GM of laboratory mice and that interactions exist between the two factors. As in other studies, these data also suggest that the CM is a more sensitive indicator of environmental effects on the gut microbiota, as compared to the FM. Specifically, while significant differences in richness were detected between treatment groups in both FM and CM, significant compositional differences were detected in only the CM. These findings reinforce the need to consider husbandry factors when comparing phenotypic data generated at different institutions or at different times, and to collect and analyze other gut regions when assessing the influence of environmental factors on the GM in general.

## Discussion

There is growing evidence that variability in husbandry practices among rodent facilities can influence rodent GM. Moreover, differences in GM have been shown to modulate model phenotypes raising the possibility that these two factors are connected: differing husbandry yield changes in GM that subsequently impact model phenotypes. Given the current concerns about reproducibility of biomedical research models (9), studies further assessing this premise are warranted. To the authors’ knowledge, this study was the first survey of changes in the GM composition due to water decontamination methods and bedding material in research mice. Results of this study, showing that water decontamination methods and bedding material can change GM composition, provide additional potential sources of GM modulation that may in turn explain why rodent phenotypes differ when experiments are performed in different housing conditions.

As demonstrated in Fig 1, rodent facilities have a large variety of methods used for water decontamination. When water acidification was introduced, several studies were performed to assess animal health and reproduction with no adverse effects demonstrated (28, 29). One of the first physiological changes associated with acidified water was decreased weight gain and water consumption (30). Another study identified changes in immune responses when animals were offered water acidified to a pH of 2.0 (31). More recently, studies have shown both changes in GM and in disease model phenotypes as a result of water acidification (16, 17, 32). In humans, while water acidification is not a commonly used practice for preventing pathogen transmission, carbonated drinks can have a pH as low as 2.0, with no known immediate direct effects on health (33).

One intriguing aspect of the present results was that even though no significant differences were detected in the overall composition of the FM, there were significant differences in richness. The changes in FM may be subtle between groups, but when the FM from arrival was compared to that of endpoint there was a significant difference in composition (S3 Fig). The significant changes in the CM composition can be an indication that cecal content is a better sample for environmental influences on the GM. The demonstrated significant main effects of both bedding and water indicate that our husbandry practices do influence the GM. The pairwise comparison (Table 2) also illustrated how different combinations of bedding and water can influence the GM.

Bedding material varies greatly between facilities, with corn cob bedding being a more popular choice, but paper based beddings are becoming increasingly common within the laboratory animal community. There are many possible reasons for corn cob to have an influence on the GM. Previous studies have demonstrated that different bedding materials do not alter the microenvironment parameters (ammonia levels, temperature, and humidity) in ventilated cages (34, 35), but there are other factors that could be influencing the changes in GM. It has been shown that mice and rats prefer alternative wood based beddings over corn cob bedding, and that corn cob can influence their sleeping habits (36–38). In a pre-diabetic mouse model, corn cob bedding reduced the efficiency of feed conversion when fed a high-fat diet (20). Corn cob bedding also contains endocrine-disrupting agents that can disrupt breeding behaviors in rats (39) and decrease aggressive behavior in the California mouse (*Peromyscus californicus*) (40). Other potential factors can be the greater amount of endotoxins and coliform levels present in corn cob bedding as compared to paper bedding (41). In a previous study evaluating the effects of changing husbandry conditions, mice switched from corn cob to a paper-based bedding demonstrated changes in microbiota composition at day 1 following the switch. However, at day 5 there were no detectable differences (21), suggesting a transient effect. While this husbandry change did not have a long-lasting effect on the FM, sampling of the cecum revealed readily detected changes.

When comparing beta diversity with PCoA, the plots with the most separation involved a group that was offered RO water (Fig 2), reflecting this treatment regimen’s capacity to influence the GM. No clear explanation can be given as to why this was the case. However, when considering the mechanistic process of reverse osmosis, it is the only decontamination method out of the four used that filters the water. The water passes through a membrane that is able to filter many compounds such as disinfectant byproducts, pesticides, endocrine-disrupting compounds, and pharmaceutical residues (42) that can potentially have a physiological influence on our rodents. This water filtration process can also filter out other compounds and minerals that would normally be available to the GM and therefore can directly influence the microbial content. Another interesting finding was that the autoclaved groups had a decreased richness of the CM when compared to all other groups. The fact that the water bottles for all groups were autoclaved in addition to other treatment in six of the groups (i.e. filtration or acidification), suggests that those additional treatments may inadvertently serve as a nidus for other bacteria or provide an environment fostering changes in the community composition. When evaluating α-diversity, significant differences were limited to the CM and were only demonstrated in the Shannon diversity index (S3 Table). The Simpson diversity index is more sensitive to abundant species (43), and therefore taxonomic units of low abundance have a smaller impact on this index. Collectively, these findings suggest that lower abundance taxa played a lesser role in the differences seen among husbandry factor combinations.

Regarding the taxa putatively susceptible to the husbandry factors under investigation, RF and two-way ANOVA identified many of the same OTUs. A total of 26 OTUs were significantly different based on two-way ANOVA with *p* values corrected to account for multiple testing, all with a significant main effect of water. Fourteen of those 26 were identified in the RF analysis as significant classifiers for the specific water and bedding combination. Several of the OTUs represented as important classifiers were OTUs that have been associated with health and disease. The genus *Akkermansia,* known to modulate the immune system and associated with metabolic diseases such as obesity (44–47), was increased in groups receiving autoclaved water. An unclassified species of *Lactobacillus* was also selected as a classifier and the relative abundance was significantly different between groups, likely due to its low relative abundance in samples from mice receiving RO-treated water. *Lactobacillus* is a genus that has gained attention through the years for its potential probiotic applications (48). In mice, certain *Lactobacillus* species have demonstrated the ability to stimulate an immunoregulatory response that allows the bacteria to persist in the bowel (49). Moreover, several *Lactobacillus* spp. have repeatedly been implicated as microbial determinants of cognitive function and behavior in mouse models (50–53) suggesting that husbandry factors affecting the GM are a critical consideration for investigators in the field of neuroscience and ethology.

In summary, water decontamination methods and bedding material used in rodent facilities can alter the GM and therefore must be considered when designing a study. Significant changes were primarily noted in cecal samples, confirming observations in previous studies (14) and suggesting that fecal sampling alone may be insufficient to unearth subtle changes in GM. Water decontamination methods vary within rodent facilities and can alter the GM composition, adding a potential variable to experimental outcomes. Accounting for and documenting these factors will aid in efforts to optimize reproducibility. It is therefore essential for investigators to provide full details as described in the ARRIVE guidelines (54) when writing a manuscript in order to increase reproducibility and ultimately translatability of our animal studies.

## Acknowledgments

The authors would like to acknowledge Karen Clifford for assistance formatting figures and the staff at the University of Missouri DNA Core and Informatics Research Core facilities.

## Author Contributions

Conceived and designed the experiments: WAB ACE CF. Performed the experiments: WAB. Analyzed the data: WAB. Contributed reagents/materials/analysis tools: ACE CF. Wrote the paper: WAB ACE CF.

## Supporting information

**S1 Table. Water decontamination methods survey.** Survey sent via e-mail through Compmed listserv on December 15, 2015.

**S1 Fig. Average relative abundance of operational taxonomic units (OTUs) in each group.** Bar graphs representing the average relative abundance of OTUs in each group for (A) fecal microbiota and (B) cecal microbiota. Each color represents a different OTU. Legend on right represents OTUs with high (>1%) relative abundance.

**S2 Table. Fecal richness and diversity analysis.** Two-way ANOVA results of richness and diversity (Shannon and Simpson diversity indices) from fecal endpoint (Day 28) samples. Considered p<0.05 significant.

**S3 Table. Cecal richness and diversity analysis.** Two-way ANOVA results of richness and diversity (Shannon and Simpson diversity indices) from cecal samples. Considered p<0.05 significant.

**S2 Fig. Hierarchical clustering of cecal samples using the most variable operational taxonomic units (OTUs).** Hierarchical clustering of the top 25 (lowest p-values corrected by false discovery rate) OTUs by one-way ANOVA of all cecal samples. Color intensity shows cube root transformed normalized abundance of OTUs in each sample. Color-coded bars at top represent bedding/water group (legend on right).

**S4 Table. Significantly different operational taxonomic units (OTUs)**. Two-way ANOVA results of significantly different (p<0.05) OTUs. Calculated using cube root transformed normalized abundance of OTUs in each sample. Numbers represent false discovery rate adjusted *p*-values.

**S3 Fig. Changes of fecal microbiota from arrival to endpoint.** Principal coordinate analysis of all fecal samples at arrival (black circles) and all different endpoint groups (legend on right). One-way PERMANOVA of ranked Bray-Curtis similarity indices results shown.

